# Disrupting ACE2 Dimerization Mitigates the Infection by SARS-COV-2

**DOI:** 10.1101/2022.04.09.487739

**Authors:** Jiaqi Zhu, Yue Su, Young Tang

## Abstract

The coronavirus disease 2019 (COVID-19) pandemic has caused over 6 million death and 460 million reported cases globally. More effective antiviral medications are needed to curb the continued spread of this disease. The infection by SARS-COV-2 virus is initiated via the interaction between the receptor binding domain (RBD) of the viral glycoprotein Spike (S protein) and the N-term peptidase domain (PD) of the angiotensin-converting enzyme 2 (ACE2) expressed on host cell membrane. ACE2 forms protein homodimer primarily through its ferredoxin-like fold domain (aka. Neck-domain). We investigated whether the dimerization of ACE2 receptor plays a role in SARS-COV-2 virus infection. We report here that the ACE2 receptor dimerization enhances the recognition of SARS-COV-2 S protein. A 43 amino acid peptide based on the N-term of Neck-domain could block the ACE2 dimerization and the interaction between RBD and ACE2, and mitigate the SARS-COV-2/host cell interaction. Our study illustrated a new route to develop potential therapeutics for the prevention and treatment of SARS-COV-2 viral infection.

## Introduction

The COVID-19 pandemic causes severe respiratory syndrome and mortality especially in adult and elderly human beings, and has paralyzed the global societal and economic development and interactions. The entry of SARS-COV-2 virus in host cells starts with the recognition of its S protein by the angiotensin-converting enzyme 2 (ACE2) on the cell membrane, which triggers the cleavage of Spike (S protein) into two subunits (S1 and S2, Fig. 1) by host cell proteases(Hoffmann et al., 2020, Shang et al., 2020), with subsequent membrane fusion between virus and host cells driven by the S2 subunit. In addition to the development of vaccines, many efforts have focused on blocking the viral entry by targeting the receptor binding domain (RBD) of S protein or PD of ACE2 with neutralizing antibodies, recombinant or soluble ACE2, peptides based on the N-term peptidase domain (PD) of ACE2 and RBD region of S protein, and small molecules targeting these domains(Datta et al., 2020, Seyedpour et al., 2020, Tay et al., 2020), with additional focuses on development of peptides and protease inhibitors blocking viral fusion and replication, and host cell endosomal and some other pathways(Seyedpour et al., 2020, Datta et al., 2020).

**Fig. 1:**
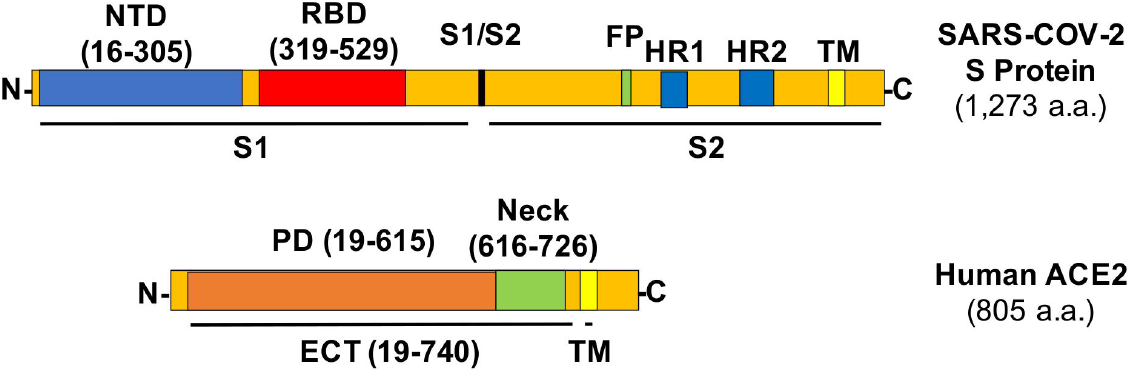
Structures of the SARS-COV-2 S Protein and Human ACE2 with Relevant Amino Acid Residues. NTD: N-term Domain. FP: Fusion Peptide. HR: Heptad Repeat. TM: Transmembrane helix.

The S protein in SARS-COV-2 viral envelop forms homotrimer with the rotation of one of the three RBDs from “down” to “up” position to be accessible by the PD of ACE2(Wrapp et al., 2020, Walls et al., 2020). The ectodomain (ECT) region of ACE2 consists of PD and the ferredoxin-like fold domain (aka. Neck-domain)(Yan et al., 2020). Crystal structure analysis revealed that ACE2 receptor naturally forms homodimer between their Neck-domains, with additional “closed” or “open” conformation developed via either formation or break of another weak interaction in PD region of the protein dimer(Yan et al., 2020). However, the significance of this dimerization for the ACE2/S protein interaction is unclear. In this study, we found that the ACE2 dimerization plays a previously unrecognized role for enhancing SARS-COV-2 infection, while inhibiting this dimerization mitigated viral infection. Our study will have important impact for the development of new therapeutics to modify the sensibility of host cells to SARS-COV-2 virus infection.

## Materials and Methods

### DNA Constructs, Cell Culture and Virus Production

For BiFC assay, the RBD domain in S protein of SARS-COV-2 virus was inserted into pBiFC-VN155 (I152L) vector, the ACE2 and its ECT/PD domains were cloned into pBiFC-VN155 (I152L) and pBiFC-VC155 vector(Kodama and Hu, 2010) (Addgene, MA, USA). The RBD, Neck-domain and different Neck-domain fragments followed directly with a stop codon were cloned into pBiFC-VN155 (I152L) vector. For protein expression, the cell membrane penetrating peptide (TAT), red fluorescence protein (DsRed) and NK-NT or NKN1 fragments were cloned into pET6xHN-N Vector (Takara, CA, USA). HEK293T cells were cultured in FP medium (DMEM containing 10% FBS, 2 mM GlutaMAX™ Supplement, 0.1 mM MEM Non-Essential Amino Acids, 50 U/mL and 50 μg/mL Penicillin-Streptomycin). For ACE2-expressing lentivirus packaging, pscALPSpuro-HsACE2 (human) (Addgene, MA, USA) were co-transfected with psPAX2 and pCMV-VSV-G packaging plasmids into HEK293T cells using FuGENE 6 (Promega, WI, USA). Supernatants containing virus were collected at 48h and 72h after transfection. For doxycycline (Dox) inducible, Spike protein pseudotyped luciferase-expressing lentivirus preparation, HEK293T cells were transfected with FUW-RLuc-T2A-PuroR(Kanarek et al., 2018) (Addgene, MA), psPAX2 and pUNO1-SARS2-S (D614G) (InvivoGen, CA) packaging plasmids using FuGENE 6. Supernatants containing virus were collected at 48h and 72h after transfection. To establish ACE2-expressed cell line (ACE2-HEK293T cells), HEK293T cells were infected with ACE2-expressing lentivirus and ACE2-positive cells were selected by 2 ug/mL of puromycin.

### Bimolecular Fluorescence Complementation (BiFC) Assay

For BiFC assay, HEK293T cells were cultured in 12-well plates and were co-transfected with 0.5 μg of each construct expressed in pBiFC-VN155 (I152L) and 0.5 μg pBiFC-VC155 vectors per well using FuGENE 6. In competition BiFC Assay, HEK293T cells were co-transfected with 0.5 μg of each construct expressed in pBiFC-VN155 (I152L) and 0.5 μg pBiFC-VC155 vectors, together with and 5 μg competitor constructs with stop codon in pBiFC-VN155 (I152L) vector. For the protein competition BiFC Assay, HEK293T cells were co-transfected with 0.25 μg of each construct expressed in pBiFC-VN155 (I152L) and 0.5 μg pBiFC-VC155 vectors in 12-well plates. FP medium containing 0.1 μg, 1 μg or 2 μg of recombinant TAT-Dsred-GFP/NK-NT/NKN1 proteins were added to the cells at 4 h after plasmid transfection. Fluorescence images were taken at 24 h and 48 h after transfection using a Nikon fluorescence microscope and fluorescence intensity was quantified by Image J.

### Protein Expression and Purification

The recombinant pET6xHN-N constructs were transformed into *Escherichia coli* strain BL21(DE3) (New England Biolabs, MA, USA). The transformed clones were cultured at 37 °C in LB medium with 100 μg/mL Carbenicillin and were induced by adding 1 mM isopropyl β-D-1-thiogalactopyranoside (IPTG) at OD 0.6-0.8 and incubated at 23 °C for overnight. For protein purification, cells were harvested by centrifugation at 5,000 rpm for 15 min and resuspended in xTractor Buffer containing DNAse I, Lysozyme solution and Protease Inhibitor Cocktail (Takara Bio USA, Inc., CA). The suspension was sonicated 10 sec for 3 times with a 30 s pause, and centrifuged at 10,000 rpm for 20 min. Supernatant was incubated with equilibrated TALON Metal Affinity Resin (Takara Bio USA, Inc., CA) for 20 min on ice. The proteins were eluted with the Elution Buffer (pH 7.0, 150 mM Imidazole, 50 mM NaH2PO4, 300 mM NaCl). And the eluted proteins were concentrated and with buffer exchanged to PBS (pH 7.4) using Protein Concentrator PES, 3K MWCO (Pierce Biotechnology, PA, USA).

### Immunoprecipitation and Western Blotting

HEK293T cells in 6-well-plate were transfected with 1 μg Myc-tagged ECT-expressing plasmids per well, and total proteins were extracted from the cells by adding Cell Lysis Buffer (Cell Signaling Technology, MA, USA) containing 500 mM NaCl, 1.5% Triton X-100 and 1 mM PMFS at 48 h after transfection. Supernatant was collected after centrifugation at 14,000 x g for 10 min and was incubated with 5 μg of recombinant GFP, NK-NT or NKN-1 proteins, and Myc-Tag (9B11) Mouse mAb (1:1000, Cell Signaling Technology, MA, USA) at 4 °C for overnight. 30 μL of protein G Agarose Beads (Cell Signaling Technology, MA, USA) slurry was added to the protein mixture and rotated at 4 °C for 3 h. Agarose beads were washed with Cell Lysis Buffer for 5 times and then resuspended in 30 μL of NuPAGE™ LDS Sample Buffer (4X) and 12 μL NuPAGE™ Sample Reducing Agent (10X) (Invitrogen, MA, USA).

Samples collected from Immunoprecipitation were separated by 10% SDS-PAGE gel and transfected on an Immobilon-P PVDF Membrane. The membrane was blocked by 5% skim milk and incubated with ACE-2 Antibody (1:2,000, Novus Biologicals, CO, USA), 6xHN Polyclonal Antibody (1:2000, Takara, CA, USA) at 4°C for overnight. The membrane was then incubated with secondary HRP-linked, Anti-rabbit IgG (1:10,000, Cell Signaling Technology, MA, USA) for 1 h at room temperature. For BiFC assay, whole proteins were extracted from HEK293T cells, separated by 10% SDS-PAGE gel and transferred onto a Immobilon-P PVDF Membrane. The membrane was incubated with ACE-2 Antibody (1:2,000, Novus Biologicals, CO, USA), Myc-Tag (9B11) Mouse mAb (1:1,000, Cell Signaling Technology, MA) and GAPDH (D16H11) XP® Rabbit mAb (1:1,000, Cell Signaling Technology, MA, USA) at 4°C overnight. The membrane was then incubated with secondary HRP-linked, Anti-rabbit IgG (1:10,000, Cell Signaling Technology, MA, USA) and Goat anti-Mouse IgG (H+L) Cross-Adsorbed Secondary Antibody, HRP (1:2,000, Thermo Fisher Scientific, MA, USA) for 1 h at room temperature. Images were developed with Clarity™ Western ECL Substrate (Bio-Rad Laboratories, CA, USA) and visualized under the ChemiDox XRS Image System (Bio-Rad Laboratories, CA, USA).

### mRNA Transfection and S Protein Pseudovirus Luciferase Assay

For mRNA preparation, GFP, NK-NT and NTN1 DNA fragments harboring T7 promoter sequence were amplified by PCR, purified, and transcribed to mRNAs *in vitro* using mMESSAGE mMACHINE™ T7 ULTRA Transcription Kit (Invitrogen, CA, USA). For the S protein pseudovirus infection and luciferase assay, ACE2-expressing HEK293T cells were seeded to 24-well plates and transfected with 2 μg of each mRNAs/well using Lipofectamine™ MessengerMAX™ Transfection Reagent (Invitrogen, CA, USA). After 24 h, cells were infected with Spike-RLuc pseudotyped lentivirus, M2rtTA lentivirus and 10 ug/mL Polybrene for 1 h. The luciferase expression was induced by adding 1 μg/mL Doxycycline at 24 h after viral infection. Luciferase activity in cells was measured using Renilla Luciferase Glow Assay Kit (Pierce Biotechnology, PA, USA) and the CLARIOstar Plus plate reader (BMG LABTECH, NC, USA) at 24 h later.

### Statistial Analysis

One way–ANOVA with Tukey’s multiple comparison post hoc test or Student’s t-test was used for data analysis. The figures were presented as mean ± standard deviation (sd). A *p*-value < 0.05 was considered statistically significant.

## Results

### Neck-Domain Is Needed for ACE2 Receptor Dimerization in Living Cells

The ACE2 ECT region consists of PD, the Neck-domain, and a long linker before the transmembrane helix (TM, excluded from ECT) (Fig. 1). The intensive polar interaction between Neck-domains is believed to primarily mediate ACE2 dimerization based on cryo-EM structural analysis(Yan et al., 2020). The bimolecular fluorescence complementation (BiFC) assay based on reconstitution of the N- and C-term fragments of the YFP protein Venus (VN and VC) has been widely applied for protein-protein interaction (PPI) study under physiological conditions(Hu et al., 2002, Kodama and Hu, 2010), allowing direct visualization of PPI in living cells with high sensitivity without exogenous reagent and complicated data process(Bhat et al., 2006, Miller et al., 2015). To verify the function of Neck-domain in living cells, we designed BiFC assays to evaluate the Neck-domain mediated ACE2 dimerization (Figs. 2A, 2B). ACE2 PD (without Neck-domain) fused with VN or VC were co-expressed in HEK293T cells. Alternatively, ACE2 ECT (with Neck-domain) fused with VN or VC were co-expressed. As expected, co-expression of PD-VN and PD-VC yielded no fluorescence, while co-expression of ECT-VN and ECT-VC produced strong fluorescence (Figs. 2C, 2D). Also, co-expression of ECT-VN with PD-VC, or ECT-VC with PD-VN did not yield any fluorescence (Figs. 2C, 2D). These data in living cells support the notion that Neck-domain is critical for ACE2 protein dimerization(Yan et al., 2020).

**Fig. 2:**
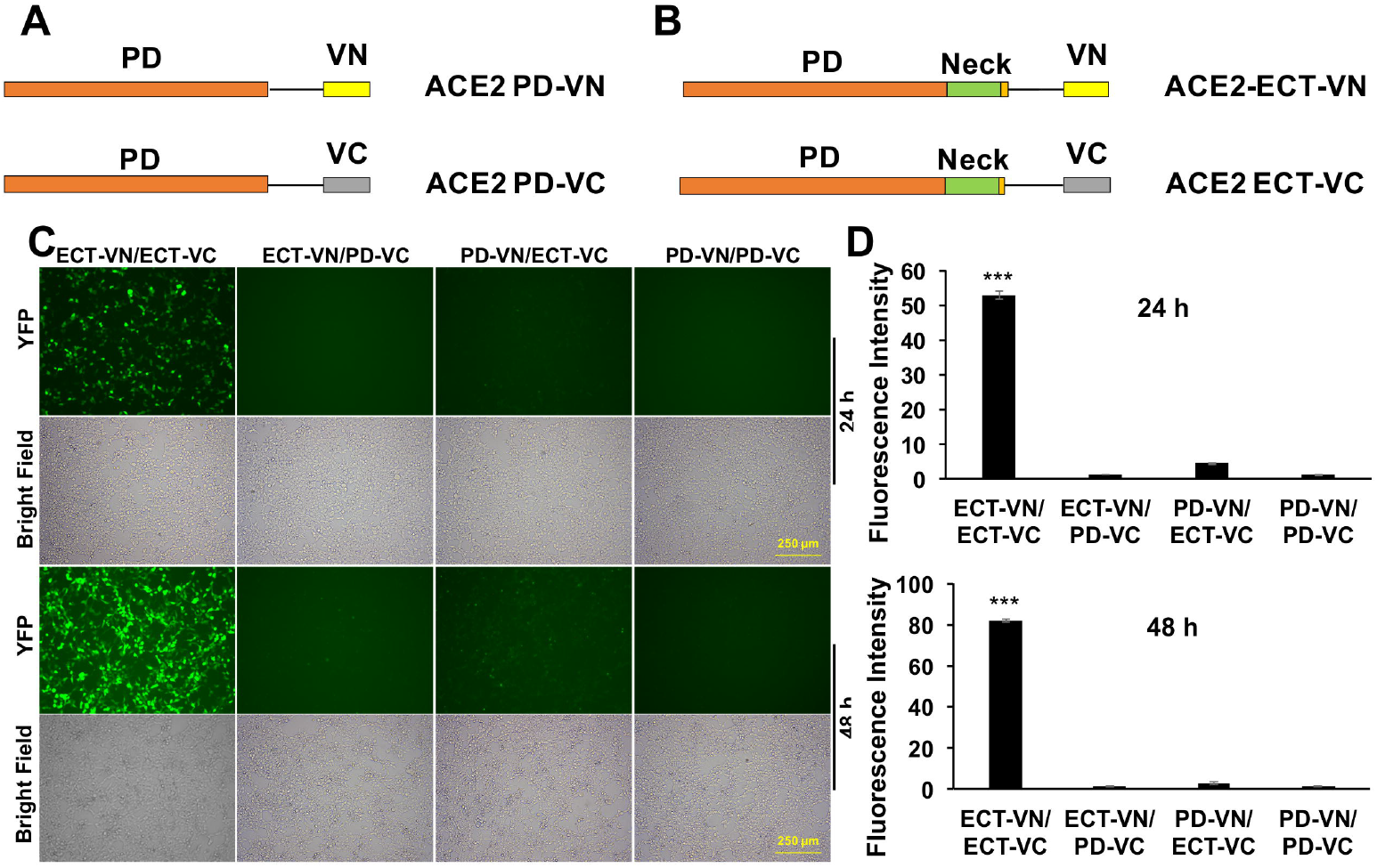
BiFC Assays to Verify the Neck-domain Function on ACE2 Dimerization. A) The PD is fused with the N-term or C-term fragment of YFP protein Venus (VN or VC). B) The ECT is fused with VN or VC. C) Representative fluorescence images of HEK293T cells co-expressing different combinations of the ACE2-ECT and ACE2-PD constructs at 24 and 48 h after the plasmid transfection. Bar = 250 µm. D) Quantitation of the fluorescence intensity at 24 (upper) and 48 h (lower) after plasmid transfection. P < 0.001, N = 3.

### Neck-Domain Is Needed for the ACE2/S Protein Interaction

To investigate the significance of Neck-domain for the PPI between human ACE2 and SARS-COV-2 S protein, we developed BiFC assay constructs by fusing S protein RBD with VN, and ACE2 PD or ECT with VC (Fig. 3A). Western blotting (WB) confirmed co-expression of S protein RBD-VN and ACE2 PD-VC or ECT-VC fusion proteins in HEK293T cells (Fig. 3B). The YFP fluorescence was then quantified at 24 h after plasmids transfection. Intriguingly, the BiFC assay revealed that ACE2 PD (w/o Neck-domain) is incapable of binding to S protein RBD in living cells, with background fluorescence signal similar as a pair of previously reported non-interactive proteins(Huang et al., 2020) - CD163 scavenger receptor cysteine-rich domain 2 (SRCR2) and porcine reproductive and respiratory virus glycoprotein GP2a (Fig. 4). On the contrary, ACE2 ECT which contains Neck-domain showed strong interaction with S protein RBD (Fig. 4). These results strongly indicate that the presence of Neck-domain is necessary for the recognition of viral S protein by ACE2. Based on the fact that Neck-domain is critical for ACE2 dimerization, our results here support the hypothesis that Neck-domain is needed for the ACE2 recognition of S protein, most likely through mediating the dimerization of ACE2.

**Fig. 3:**
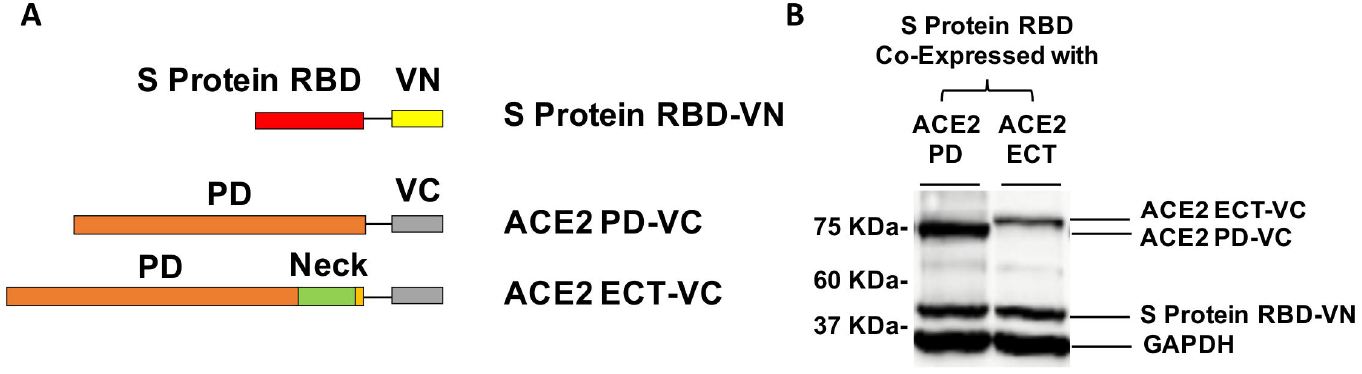
Design of BiFC Assay to Detect the ACE2/S Protein PPI. A) The RBD of S protein was fused with VN, and PD (a.a. 19-615) or ECT (a.a. 19-740) of ACE2 was fused with VC. B) Western Blot detection of the recombinant proteins co-expressed in HEK293T cells for BiFC assay using specific antibodies. GAPDH served as the loading control.

**Fig. 4:**
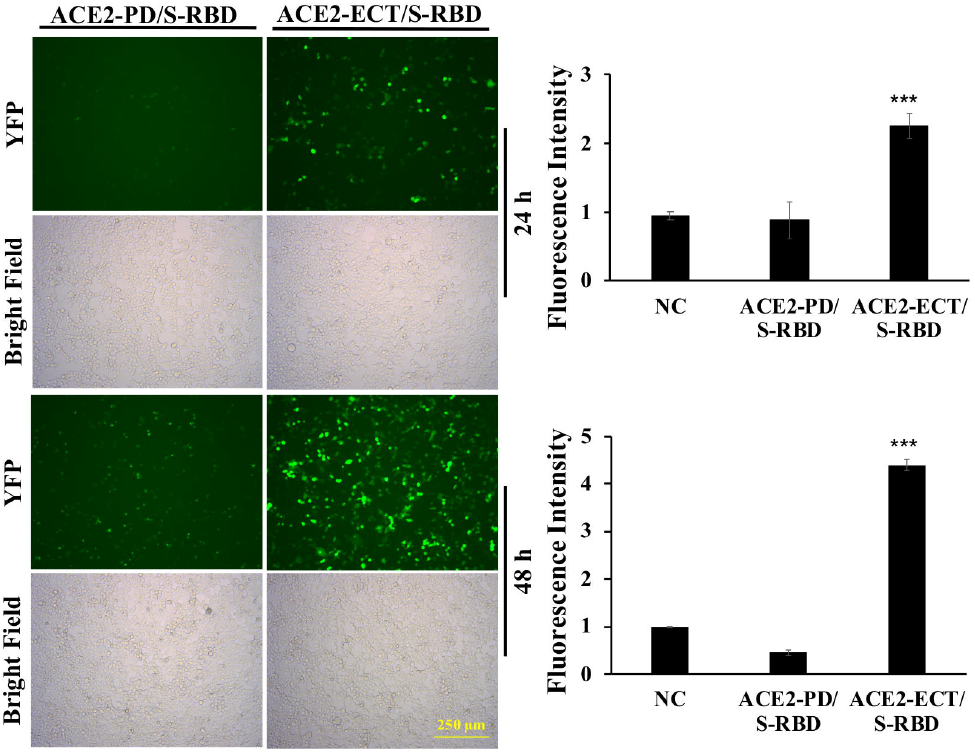
ACE2 Neck-domain Is Critical for ACE2/S Protein Interaction. Left: Representative BiFC fluorescence images of HEK293T cells co-transfected with ACE2-PD-VC or ACE2-ECT-VC and S protein RBD-VN plasmids at 24 and 48 h, respectively. Bar = 250 µm. Right: Quantitation of the relative fluorescence intensity. p < 0.001, N = 3. NC: negative control using a pair of non-interactive proteins (CD163-SRCR2 and GP2a) as described in the main text.

### Screening Parts of Neck-Domain Critical for ACE2 Dimerization and S protein Recognition

To investigate which part of Neck-domain is critical for ACE2 dimerization, we designed a competition assay by co-expressing different segments of Neck-domain as the competitor (Fig. 5A) in the ECT-VN/ECT-VC BiFC system (Fig. 2B). The competitor plasmids were added at 5-fold more than ECT-VN/ECT-VC vectors, with data collected at 72 h after plasmid transfection to allow the proper expression of ECT-VN/ECT-VC vectors. As expected, PD without Neck-domain failed to interfere with the interaction between ECT-VN and ECT-VC (Fig. 5B). The competition BiFC screening revealed that N-term residues 616-671 of Neck-domain (NK-NT, Fig. 5A) strongly inhibited the PPI between ECT-VN and ECT-VC (Fig. 5B). Interestingly, overexpression of NK-MT caused aggregation and death of cells (Fig. 5B). To investigate how the disturbance of ECT dimerization could impact the PPI between S protein RBD and ECT, we co-expressed these segments of Neck-domain as competitors in RBD-VN/ECT-VC BiFC system (Fig. 3A). Over-expression of PD as a competitor could only slightly inhibit the recognition of RBD by ECT (Fig. 5C). However, NK-NT fragment exhibited the most significant inhibitory effect on the PPI between RBD and ECT (Figs. 5C, 5D), similar as in the ECT dimerization BiFC assay (Fig. 5B). Further screening of different parts of NK-NT identified that residues 616-658 of Neck-domain (NKN1, Fig. 5A) shares similar potency as NK-NT in blocking the RBD/ECT interaction (Fig. 5E). This correlates well with the comparable efficiency between NK-NT and NKN1 on the inhibition of full length ACE2 protein dimerization in BiFC assay (Fig. 5F).

**Fig. 5:**
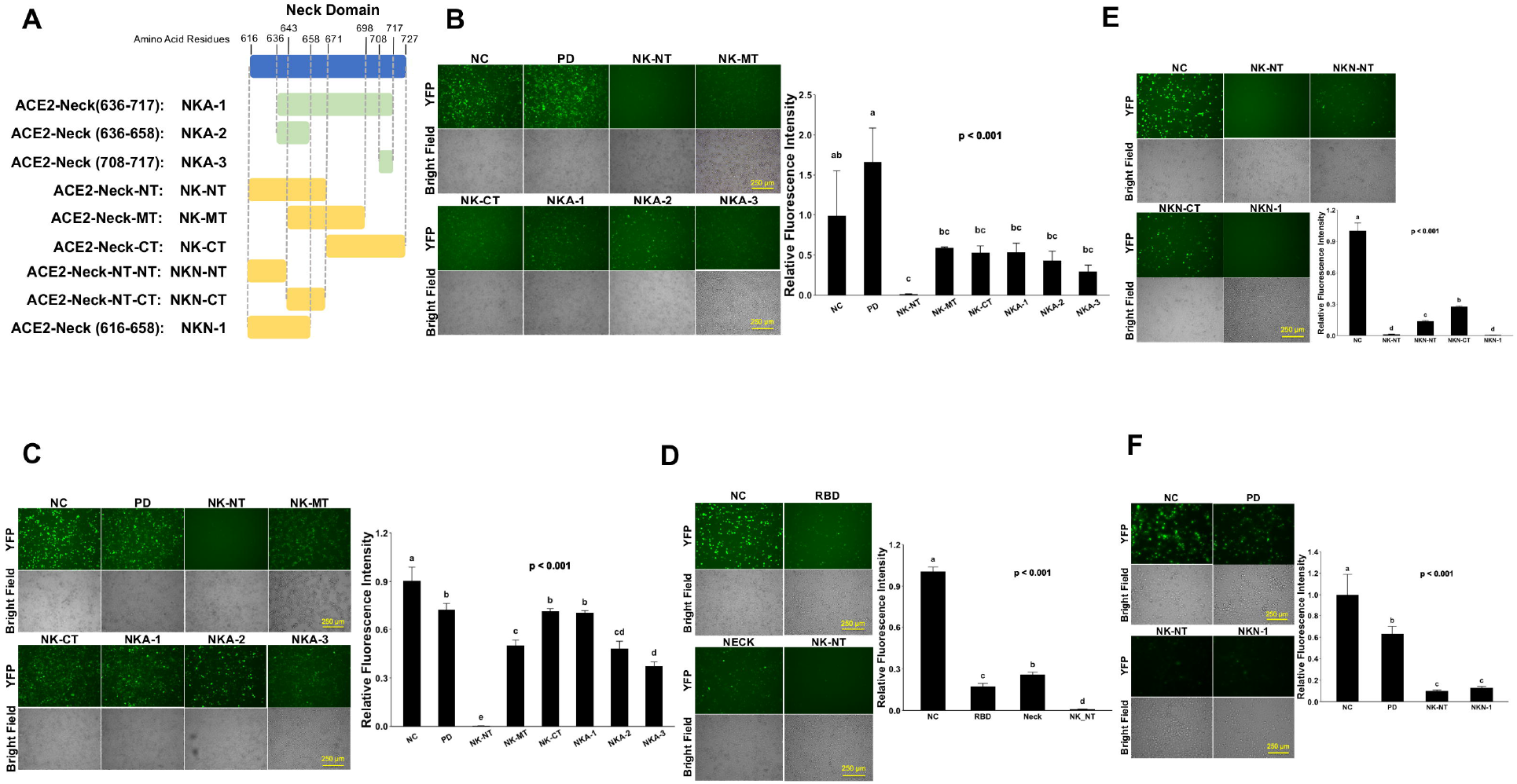
Screening Parts of Neck-domain Critical for ACE2 Dimerization and Recognition of S Protein. A) Schematic diagram of different Neck-domain fragments for BiFC screening. B) Competition ECT-VN/ECT-VC BiFC results with the co-expression of different Neck-domain fragments as competitors. Left: Representative BiFC fluorescence images. Bar = 250 µm. Right: Quantitation of the relative fluorescence intensity. p <0.01, N = 3. NC: Negative control of ECT-VN/ECT-VC BiFC with no competitors. C) Competition RBD-VN/ECT-VC BiFC results with the co-expression of different Neck domain fragments as competitors. Left: Representative BiFC fluorescence images. Bar = 250 µm. Right: Quantitation of the relative fluorescence intensity. p <0.001, N = 3. NC: Negative control of RBD-VN/ECT-VC BiFC with no competitors. D) Competition RBD-VN/ECT-VC BiFC results with RBD, Neck-domain, and NK-NT as competitors. Left: Representative BiFC fluorescence images. Bar = 250 µm. Right: Quantitation of the relative fluorescence intensity. p <0.001, N = 3. E) Comparing competition RBD-VN/ECT-VC BiFC results with additional fragments of NK-NT as competitors. Left: Representative BiFC fluorescence images. Bar = 250 µm. Right: Quantitation of the relative fluorescence intensity. p <0.001, N = 3. F) Comparing competition full length ACE2-VN/ACE2-VC BiFC results with PD, NK-NT, and NKN1 as competitors. Left: Representative BiFC fluorescence images. Bar = 250 µm. Right: Quantitation of the relative fluorescence intensity. p <0.001, N = 3.

In order to double verify the results of the competition BiFC assays, we engineered histidine and asparagine (HN)-tagged protein constructs with fusion of cell penetrating peptide TAT(Frankel and Pabo, 1988), red fluorescence protein DsRed, and NK-NT (Fig. 6A, Upper). The TAT-DsRed(Tang et al., 2011) and TAT-DsRed-NK-NT recombinant proteins were purified and quantified, and added directly to culture medium of 293T cells transfected with ECT-VN/ECT-VC or ACE2-VN/ACE2-VC plasmids (Fig. 6A, Lower). Compared with the TAT-DsRed control, one μg/mL and two μg/mL TAT-DsRed-NK-NT significantly, though not completely inhibited the dimerization between ECT (Fig. 6B) or ACE2 (Fig. 6C). We also expressed HN-tagged DsRed-NKN1, DsRed-NK-NT fusion proteins (Fig. 6D) and conducted Co-immunoprecipitation (Co-IP) experiment using total lysates of 293T cells over-expressing Myc-tagged ECT. We found that both NKN1 and NK-NT recombinant proteins interacted with ECT (Fig. 6E). Taken together, our results revealed a 43-a.a. peptide (NKN1) as the critical segment in Neck-domain necessary for the PPI between ACE2/S protein by mediating ACE2 dimerization.

**Fig. 6:**
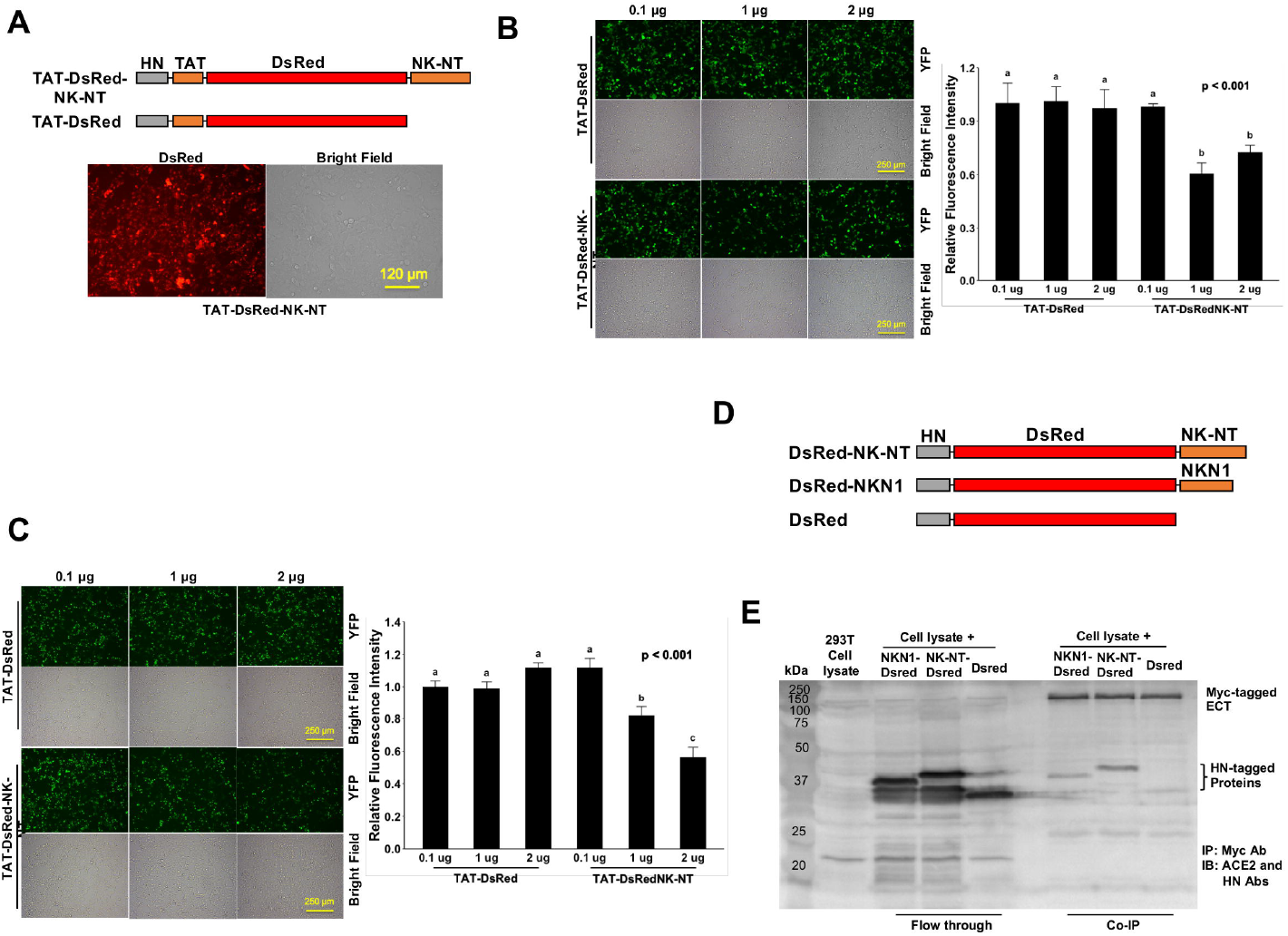
Recombinant NK-NT and NKN1 Proteins Bind to ECT and Block ACE2 Dimerization. A) Upper: Schematic diagram of histidine and asparagine (HN)-tagged recombinant protein constructs with cell penetrating peptide TAT. Lower: Representative image showing the HN-TAT-NK-NT recombinant protein transfected into 293T cells. Bar = 120 µm. B) Competition ECT-VN/ECT-VC BiFC results with the TAT-DsRed control or TAT-DsRed-NK-NT as competitors. Left: Representative BiFC assay fluorescence images. Bar = 250 µm. Right: Quantitation of the relative fluorescence intensity. p <0.001, N = 3. C) Competition ACE2-VN/ACE2-VC BiFC assay results with the TAT-DsRed control or TAT-DsRed-NK-NT as competitors. Left: Representative BiFC fluorescence images. Bar = 250 µm. Right: Quantitation of the relative fluorescence intensity. p <0.001, N = 3. D) Schematic diagram of HN-tagged recombinant NK-NT and NKN1 protein constructs. E) Co-IP results of purified HN-tagged recombinant DsRed, NK-NT and NKN1 proteins incubated with lysates of the 293T cells expressing Myc-tagged ECT.

### NK-NT and NKN1 mRNAs Inhibit SARS-COV-2 Viral Infection

Previous cryl-EM structure study mapped extensive polar interactions for ACE2 dimerization in the second (a.a. 636-658, NKA-2) and fourth (a.a. 708-717, NKA-3) helices of Neck-domain(Yan et al., 2020) (Fig. 7A), which may explain the partial activities we observed for NKA-2 and NKA-3 segments in blocking ECT dimerization (Fig. 5B). However, NK-NT and NKN1 fragments exhibited the greatest potency in blocking ECT dimerization (Fig. 5B), indicating that in addition to the second helix, additional residue(s) before it (residues 616-636) also has critical function for ACE2 dimerization (Fig. 7A). One such could be Y633 that forms cation-π interaction with R710 in the fourth helix(Yan et al., 2020) (Fig. 7A). In order to test whether the expression of NK-NT and NKN1 fragments can interfere with the SARS-COV-2 viral infection, we in vitro transcribed and purified the NK-NT and NKN1 mRNAs with the GFP mRNA as negative control (Fig. 7B). Two μg of mRNA per well were transfected to 293T cells stably expressing human ACE2 in 24-well plates. After 24 h of transfection, the cells were infected with Renilla luciferase-expressing FUW-lentivirus pseudotyped with SARS-COV-2 S protein (D614G) as the envelope protein, and the luciferase activity were evaluated at 48 h after viral infection. We found that compared with the GFP mRNA, the transfection of both NK-NT and NKN1 mRNAs significantly inhibited S protein pseudotyped lentivirus infection of the cells (Fig. 7C).

**Fig. 7:**
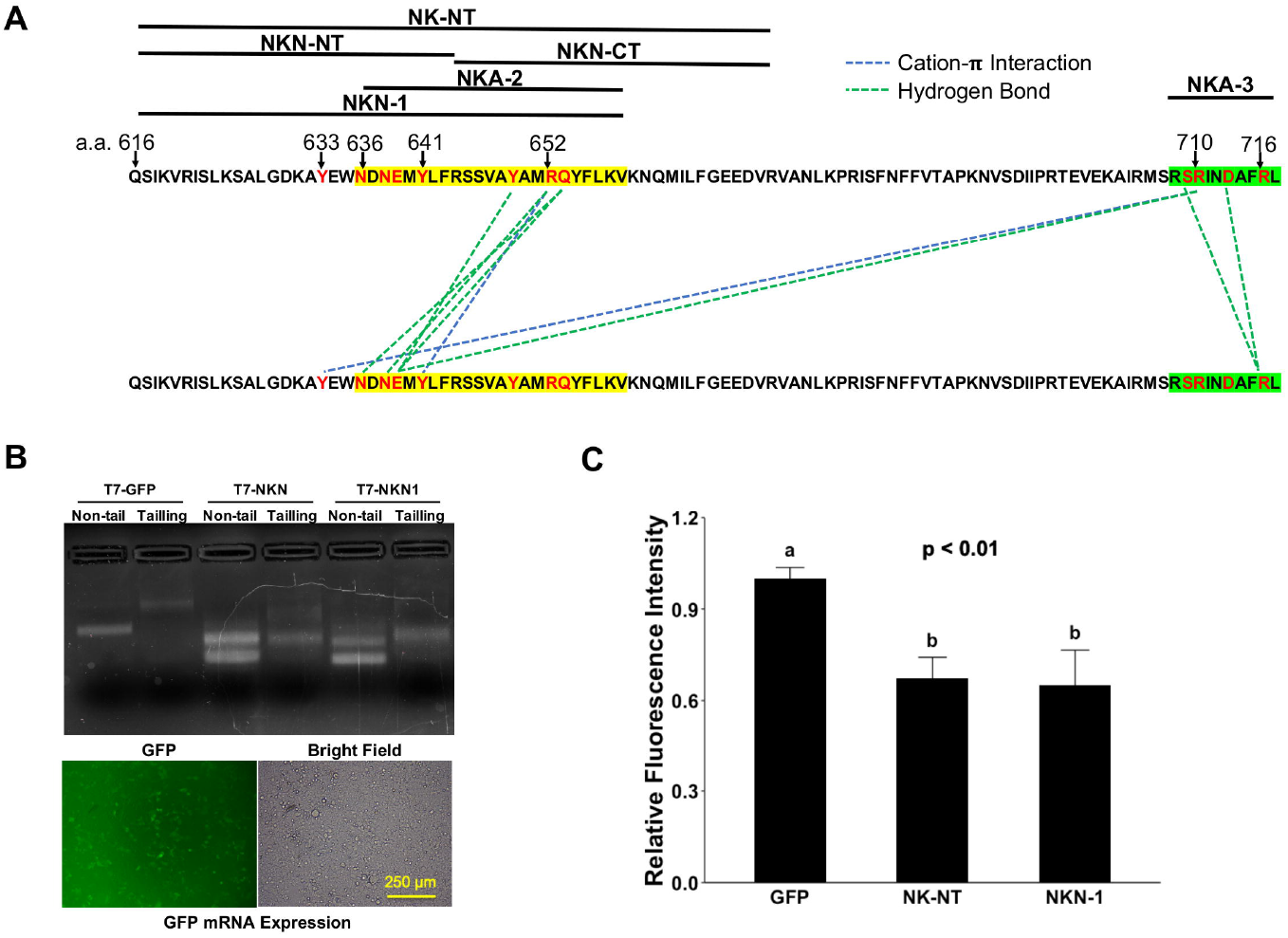
NK-NT and NKN1 mRNAs Mitigate SARS-COV-2 Pseudovirus Infection. A) Residues in NK-NT, NKN1, and Neck-domain critical for ACE2 dimerization. The second and forth helices are highlighted by yellow and green, respectively. B) Upper: GFP, NK-NT and NKN1 mRNAs transcribed in vitro via DNA constructs with a T7 promoter. Non-tail: Purified mRNAs transcribed in vitro. Tailing: Purified mRNAs with addition of poly-A tail. Lower: Expression of GFP mRNA after transfection into 293T cells. Bar = 250 µm. C) Relative luminescence intensity in 293T cells in 24-well plates transfected with 2 µg of GFP, NK-NT or NKN1 mRNAs per well and followed by infection of S protein pseudotyoped lentivirus expressing Renilla luciferase. p<0.01, N = 3.

## Discussion

The COVID-19 pandemic has caused nearly 500 million infection cases and over 6 million death globally. Numerous research has been done to better understand the mechanism of the SARS-COV-2 viral infection and its pathological effect in order to combat this disease. In this study we tried to understand the fundamental aspects that determine this virus/host cell interaction to aid in the design of effective therapeutics, and used uniquely designed serial BiFC assays to verify the essential role of Neck-domain for the ACE2/S protein PPI. We found that only the ACE2 fragment capable of self-dimerization can effectively bind to S protein of the SARS-COV-2. This is consistent with the previous observation from cryl-EM structural study that the interaction with S protein RBD was only observed in ACE2 dimer with “closed” conformation(Yan et al., 2020). These data indicate that conformational changes caused by dimerization could affect the binding affinity of ACE2 to S protein. We further identified that the presence of Neck-domain is critical for the successful ACE2/S protein recognition. This indicates that the disruption of Neck-domain mediated ACE2 dimerization in host cells might have significant impact on S protein recognition.

We further identified the minimal peptide sequence from residues 616-658 (NKN1) in Neck-domain is necessary for blocking ACE2 dimerization and the PPI between ACE2 and S protein. It is interesting that NKA-1 fragment (Fig. 5A) containing both the second and forth helices could not effectively inhibit the PPI between ACE2 or ACE2/S protein. One reason could be the potential self-association of NKA-1 fragments caused by extensive hydrogen bond and Cation-π interaction among residues within these two helices (Fig. 7A). By expressing NKN1 or NK-NT peptides in ACE2-expressing cells, we confirmed that these peptides could inhibit the S protein pseudotyped lentivirus infection. These results correlate well with a previous study showing that artificial dimerization of ACE2 PD by fusion with the mouse IgG Fc has much lower efficiency than that of ACE2-ECT in blocking SARS-COV-2 pseudovirus infection(Li et al., 2020). Our study lets us to predict that in host cell surface, ACE2 receptors form homodimer to enable S protein recognition, and the dimerization mediated by Neck-domain is necessary for ACE2/S protein interaction and the SARS-COV-2 viral infection. Thus, drugs interfering with Neck-domain to disrupt ACE2 homodimerization would have the potential to minimize the susceptibility of host cells to SARS-COV-2 virus.

We therefore expect that future research would identify refined protein peptides based on Neck-domain and small molecules that can effectively block the ACE2 dimerization, to enhance the host cell “immunity” to SARS-COV-2 virus. These could be applied to either protect the host from viral infection or curb subsequent viral amplification and viremia in infected hosts. These drugs could also have extended effect on preventing the infection by other members of the coronavirus family that utilize the recognition of ACE2 receptor to enter the cells.

## Data Availability Statement

All data presented in the study are included in the article, further inquiries can be directed to the corresponding author.

## Author Contributions

YT and JZ conceived and designed the study and wrote the manuscript. JZ and YS performed the experiments, conducted data analysis, and revision of the manuscript.

## Acknowledgement

This study was supported by the University of Connecticut OVPR COVID-19 Rapid Start Funding.

## Confliction of Interest

The authors declare that they have no conflict of interest.

## Notes

### Competing Interest Statement

The authors have declared no competing interest.

### Summary of Updates

typos, grammar issues in the main text and figure legends

## References

Bhat, R. A., Lahaye, T. & Panstruga, R. 2006. The visible touch: in planta visualization of protein-protein interactions by fluorophore-based methods. Plant Methods, 2, 12.

Datta, P. K., Liu, F., Fischer, T., Rappaport, J. & Qin, X. 2020. SARS-CoV-2 pandemic and research gaps: Understanding SARS-CoV-2 interaction with the ACE2 receptor and implications for therapy. Theranostics, 10, 7448–7464.

Frankel, A. D. & Pabo, C. O. 1988. Cellular uptake of the tat protein from human immunodeficiency virus. Cell, 55, 1189–93.

Hoffmann, M., Kleine-Weber, H., Schroeder, S., Kruger, N., Herrler, T., Erichsen, S., Schiergens, T. S., Herrler, G., Wu, N. H., Nitsche, A., Muller, M. A., Drosten, C. & Pohlmann, S. 2020. SARS-CoV-2 Cell Entry Depends on ACE2 and TMPRSS2 and Is Blocked by a Clinically Proven Protease Inhibitor. Cell, 181, 271–280 e8.

Hu, C. D., Chinenov, Y. & Kerppola, T. K. 2002. Visualization of interactions among bZIP and Rel family proteins in living cells using bimolecular fluorescence complementation. Mol Cell, 9, 789–98.

Huang, C., Bernard, D., Zhu, J., Dash, R. C., Chu, A., Knupp, A., Hakey, A., Hadden, M. K., Garmendia, A. & Tang, Y. 2020. Small molecules block the interaction between porcine reproductive and respiratory syndrome virus and CD163 receptor and the infection of pig cells. Virol J, 17, 116.

Kanarek, N., Keys, H. R., Cantor, J. R., Lewis, C. A., Chan, S. H., Kunchok, T., Abu-Remaileh, M., Freinkman, E., Schweitzer, L. D. & Sabatini, D. M. 2018. Histidine catabolism is a major determinant of methotrexate sensitivity. Nature, 559, 632–636.

Kodama, Y. & Hu, C. D. 2010. An improved bimolecular fluorescence complementation assay with a high signal-to-noise ratio. Biotechniques, 49, 793–805.

Li, Y., Wang, H., Tang, X., Fang, S., Ma, D., Du, C., Wang, Y., Pan, H., Yao, W., Zhang, R., Zou, X., Zheng, J., Xu, L., Farzan, M. & Zhong, G. 2020. SARS-CoV-2 and three related coronaviruses utilize multiple ACE2 orthologs and are potently blocked by an improved ACE2-Ig. J Virol.

Miller, K. E., Kim, Y., Huh, W. K. & Park, H. O. 2015. Bimolecular Fluorescence Complementation (BiFC) Analysis: Advances and Recent Applications for Genome-Wide Interaction Studies. J Mol Biol, 427, 2039–2055.

Seyedpour, S., Khodaei, B., Loghman, A. H., Seyedpour, N., Kisomi, M. F., Balibegloo, M., Nezamabadi, S. S., Gholami, B., Saghazadeh, A. & Rezaei, N. 2020. Targeted therapy strategies against SARS-CoV-2 cell entry mechanisms: A systematic review of in vitro and in vivo studies. J Cell Physiol.

Shang, J., Ye, G., Shi, K., Wan, Y., Luo, C., Aihara, H., Geng, Q., Auerbach, A. & Li, F. 2020. Structural basis of receptor recognition by SARS-CoV-2. Nature, 581, 221–224.

Tang, Y., Lin, C. J. & Tian, X. C. 2011. Functionality and transduction condition evaluation of recombinant Klf4 for improved reprogramming of iPS cells. Cell Reprogram, 13, 99–112.

Tay, M. Z., Poh, C. M., Renia, L., Macary, P. A. & Ng, L. F. P. 2020. The trinity of COVID-19: immunity, inflammation and intervention. Nat Rev Immunol, 20, 363–374.

Walls, A. C., Park, Y. J., Tortorici, M. A., Wall, A., Mcguire, A. T. & Veesler, D. 2020. Structure, Function, and Antigenicity of the SARS-CoV-2 Spike Glycoprotein. Cell, 181, 281–292 e6.

Wrapp, D., Wang, N., Corbett, K. S., Goldsmith, J. A., Hsieh, C. L., Abiona, O., Graham, B. S. & Mclellan, J. S. 2020. Cryo-EM structure of the 2019-nCoV spike in the prefusion conformation. Science, 367, 1260–1263.

Yan, R., Zhang, Y., Li, Y., Xia, L., Guo, Y. & Zhou, Q. 2020. Structural basis for the recognition of SARS-CoV-2 by full-length human ACE2. Science, 367, 1444–1448.

